# Optimal Maternal Feeding Isotopic Niche: influence of breeder trophic behaviour on larval growth and survival in bluefin tuna species

**DOI:** 10.1101/2025.09.03.673896

**Authors:** José M Quintanilla, Ricardo Borrego-Santos, Estrella Malca, Isabel Riveiro, Francisco J. Abascal, Miquel Planas, Rasmus Swalethorp, Michael R. Landry, Raúl Laiz-Carrión

## Abstract

Maternal effects play a fundamental role in shaping early larval growth and survival in marine fishes. This study explores the relationship between maternal trophic ecology and larval growth in bluefin tunas, with a focus on Southern Bluefin Tuna (SBT) and an expanded dataset from multiple Atlantic Bluefin Tuna populations incorporated into the General Bluefin Model (GBM). Daily growth and stable isotopes (δ¹⁵N and δ¹³C) were obtained from 80 pre-flexion SBT larvae and 355 pre-flexion larvae from the GBM dataset. Results revealed a significant negative linear trend between larval age and δ¹⁵N values, consistent with the gradual attenuation of maternally inherited isotopic signatures during development. Faster growing larvae had higher growth rates showed significantly lower δ¹⁵N and δ¹³C values, indicating that maternal trophic behaviour and their prey sources critically influence larval growth potential. Maternal isotopic niche breadth, inferred from larval isotope data, was markedly narrower in groups with optimal larval growth, suggesting that specialized (stenophagous) maternal feeding strategies promote enhanced offspring performance. These patterns were observed consistently in two bluefin species across seven different populations, despite geographic and temporal variability, highlighting a robust ecological link between maternal foraging behaviour and larval development. From these findings, we introduce the hypothesis of an Optimal Maternal Feeding Isotopic Niche, representing a constrained isotopic range associated with increased larval growth and survival. This framework advances our understanding of the influence of maternal trophic ecology on offspring fitness and offers valuable insights for the conservation and management of highly migratory pelagic species with complex reproductive strategies.

## 1. Introduction

Maternal effects play a pivotal role in shaping the early development, survival, and quality of fish larvae. These effects stem from various maternal contributions, including the transfer of hormones, nutrients, immune factors, and environmental conditions to the eggs, all of which can significantly influence larval phenotype and fitness (Lam, 1994; McCormick, 1999; Swain and Nayak, 2009).

Among these contributions, the maternal transfer of nutrients and immune factors is particularly critical for larval health. The quality and quantity of such maternal provisions depend on the mother’s physiological condition, age, weight, and environmental history (Conover and Schultz, 1997; Swain and Nayak, 2009; Fan et al., 2019; Hiraoka et al., 2019; Rahman et al., 2021). The diet prior to spawning has been shown to affect reproductive traits such as gonadosomatic index (Belgrad and Griffen, 2016), hatch size (Reznick and Yang, 1993; Marshall et al., 2008), and especially the biochemical composition of eggs (Yoshida et al., 2011; Schlotz et al., 2013). Maternal provisioning not only determines the initial size and quality of eggs and, hence, larval growth (Garrido et al., 2015; Fennie et al. 2023, Walsh et al. 2024) but also shapes the biochemical and isotopic composition of larvae during their most vulnerable early stages (Uriarte et al., 2016; Ohshimo et al., 2018).

Maternal isotopic transmission has been traced in perciforms to offspring (Starrs et al., 2014), and these maternal effects remain detectable until key developmental milestones, such as notochord flexion, after which larval isotopic values begin to reflect exogenous feeding (Uriarte et al., 2016). Consequently, maternal isotopic effects are most pronounced in pre-flexion stages and provide valuable markers for distinguishing pre- and post-flexion stages and for studying early trophic ecology (García et al., 2017; Laiz-Carrión et al., 2019).

Several authors have reported that stable isotope values are influenced by growth rates (Suzuki et al., 2005; Watanabe et al., 2005) and that a close relationship exists between feeding behaviour, as inferred from SIA, and growth potential during early life stages (Laiz-Carrión et al., 2011, 2013, this issue; Quintanilla et al., 2015, 2020).

In tuna species characterized by high energetic demands on development, maternal effects include the transfer of nutrients, energy reserves, and stable isotopic signatures (δ¹⁵N and δ¹³C) from mother to offspring. Recent studies of Atlantic Bluefin Tuna (ABT) have shown that maternal isotopic inheritance is closely linked to larval growth potential and maternal energy investment. Notably, larvae exhibiting optimal growth have lower δ¹⁵N values and higher energy allocation toward somatic development (Quintanilla et al., 2023, 2024). Furthermore, interannual and regional variability of maternal trophic niches and environmental conditions have been shown to influence larval growth and survival, highlighting the relevance of maternal effects for understanding population dynamics and informing fisheries management (Ohshimo et al., 2018; Quintanilla et al., 2023, 2024).

The Southern Bluefin Tuna (*Thunnus maccoyii*, SBT) is a highly migratory and commercially important species inhabiting temperate waters of the Southern Hemisphere, from the eastern Atlantic to the Indian and southwestern Pacific Oceans, between 30°S and 50°S (Caton, 1991). Unlike the closely related Atlantic Bluefin Tuna (ABT), the reproductive behaviours and spawning habitats of SBT remain poorly understood. Mature SBT migrate to a single known spawning area in the northeastern Indian Ocean (IO), between Java and Australia (Farley and Davis, 1998; Patterson et al., 2008). However, regular larval sampling is lacking, and most available data on early life ecology stem from surveys conducted decades ago. These early surveys (from the mid-1950s to early 1980s) identified a primary spawning zone approximately bounded by 7–20°S and 110–125°E (Nishikawa et al., 1985).

Understanding the mechanisms and consequences of maternal effects in bluefin tuna species is essential for predicting larval performance, recruitment success, and population resilience under changing environmental conditions. From this perspective, the present study examines somatic variables alongside individual δ¹⁵N and δ¹³C during the pre-flexion-stage SBT larvae when maternal inheritance is most pronounced. We assessed the influence of maternally inherited trophic markers on early larval growth and survival to infer female feeding strategies from maternal isotopic signatures (δ¹⁵N_maternal_, δ¹³C_maternal_).

In addition, we applied a similar analytical framework to a broader dataset designated as the “General Bluefin Model” (hereafter GBM) that includes preflexion-stage SBT larvae from the IO population together with Atlantic Bluefin Tuna (ABT) populations over several years and in different spawning grounds. This broader comparison allowed us to test the model’s robustness and assess whether our findings extend across bluefin tuna species, while introducing a new hypothesis of an “Optimal Maternal Feeding Isotopic Niche” (OMFIN) to explain how maternal isotope signatures influence larval growth potential

## 2. Material and methods

### 2.1. Ichthyoplankton collections and sample processing

#### 2.1.1. Southern Bluefin Tuna

As part of the 2nd International Indian Ocean Expedition (IIOE-2), larvae and eggs of SBT, *Thunnus maccoyii* (Castelnau, 1872) along with environmental data, were collected during the BLOOFINZ-IO oceanographic cruise aboard R/V *Roger Revelle*. The expedition was conducted from 20 January to 14 March 2022, coinciding with the peak spawning season of SBT at its only known spawning ground in the eastern Indian Ocean **(Fig. 1)**.

**Figure 1.**
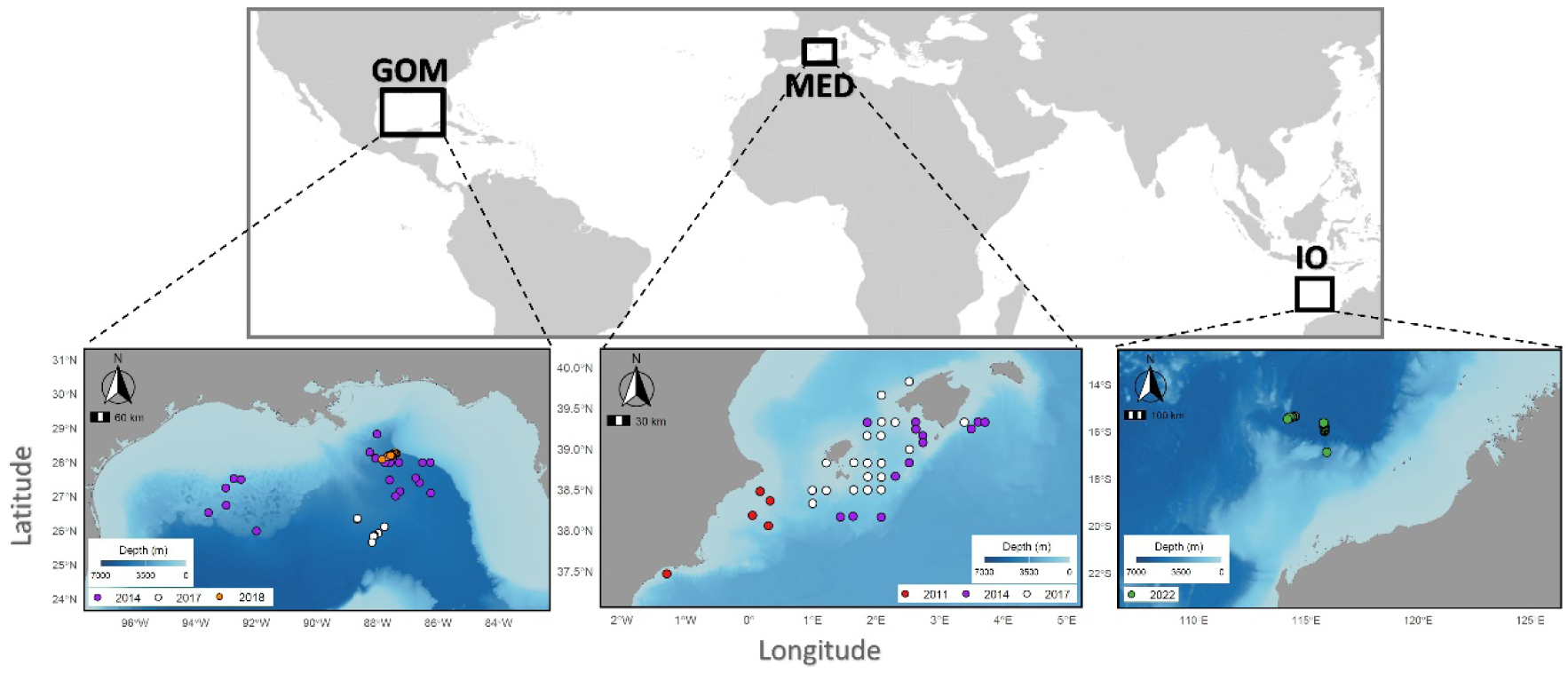
Geographical location of bluefin tuna larval sampling included in this study from the Gulf of Mexico (GOM), Nothwestern Mediterranean Sea (MED) and eastern Indian Ocean (IO).

Ichthyoplankton samples were collected using double oblique tows from 25 m depth to the surface with a 90-cm Bongo net (500 and 1000-µm mesh) and a 1-m² surface net (1000-µm mesh) (Table 1). Upon detection of Southern bluefin tuna (SBT) larval aggregations of sufficient density (≥ 5 individuals on one side of a standard oblique bongo tow), the corresponding water parcel was first geolocated using a satellite-tracked surface drifter. Intensive sampling was subsequently initiated within 1 nm of the drifter at ∼3–6 h intervals, following the drifter’s trajectory over a continuous 3–4 days period. A total of four such Lagrangian sampling experiments—designated Cycles 1 through 4—were conducted, during which high-resolution measurements of hydrographic structure, biogeochemical variables, and planktonic rate processes were carried out. Additional larval sampling was performed at discrete stations during transit legs between cycles, as well as along basin-scale transects spanning the Argo Basin at both the outset and conclusion of the cruise. For overall study design details, see Landry et al. (this issue).

**Table 1.** Collection details for Southern Bluefin Tuna (*Thunnus maccoyii*) in the Indian Ocean (IO) and Atlantic Bluefin Tuna (*Thunnus thynnus*) larvae in the Gulf of Mexico (GOM), the Mediterranean Sea (MED)).

| Spawning ground and survey year | Sampling gear and mesh size | Number of larvae | Standard length (mm) |
| --- | --- | --- | --- |
| **Indian Ocean** |  |  |  |
| IO22 | Bongo 90 - 500,1000 μm / Neuston net - 1000 μm | 80 | 2,97 - 6,34 |
| **Gulf of Mexico** |  |  |  |
| GOM14 | S10 - 500 μm | 54 | 4,22 - 5,84 |
| GOM17 | Bongo 90 - 500 μm | 49 | 4,28 - 7,06 |
| GOM18 | Bongo 90 - 500 $\mu\text{m}$ | 36 | 4,10 - 6,36 |
| Subtotal | — | 139 | — |
| **Northwest Mediterranean** |  |  |  |
| MED11 | Bongo 60 - 300 $\mu\text{m}$ / Bongo 90 - 1000$\mu\text{m}$ | 27 | 3,77 - 5,59 |
| MED14 | Bongo 90 - 500 $\mu\text{m}$ | 65 | 2,41 - 6,69 |
| MED17 | Bongo 90 - 500 $\mu\text{m}$ | 44 | 3,18 - 5,82 |
| Subtotal | — | 136 | — |
| **TOTAL** |  | **355** | **2,41 -7,06** |

SBT larvae were identified shipboard based on morphological, meristic, and pigmentation features following Nishikawa et al. (1985, 1987), and subsequently preserved at -80 °C for further genetic, isotopic, and ageing analyses at the Oceanographic Center of Málaga (IEO-CSIC, Spain). DNA was extracted from each larva for post-cruise species verification via multiplex PCR, distinguishing SBT from congener (*Thunnus albacares*, *T. alalunga*, *T. obesus*) and tuna-like species (*Katsuwonus pelamis*) present in the sampling area (see Malca et al., this issue for details). In order to detect the maternal isotopic influence, only preflexion-stage SBT larvae were selected for this study (see Borrego-Santos et al. and Laiz-Carrión et al. this issue for the complementary post-flexion larval).

#### 2.1.2. Atlantic Bluefin Tuna

ABT (*Thunnus thynnus* (Linnaeus, 1758)) larvae included in the GBM dataset together with SBT population were collected across several years from six surveys (Table 1) conducted in the two primary spawning areas of the species: the Gulf of Mexico (GOM) and the Northwestern Mediterranean Sea (MED) (Fig. 1). Collection procedures including net types (Bongo (90 and 60-cm), mesh size (500 and 1000-µm), depths sampled (0-30 m), duration of tows (∼10 min), speed of vessels (∼2-3 knots) were similar to those used for SBT and are detailed in García et al. 2012, Malca et al. (2022, 2023) and Quintanilla et al. (2023, 2024) (Table 1).

### 2.2. Larval standard length and dry weight

For daily growth estimation, we exclusively used preflexion stage larvae with notochord flexion assessed from digitized, calibrated images (see Borrego-Santos et al., this issue). In the laboratory, larval standard length (SL, mm) was measured using calibrated images from ImageJ 1.44a (NIH, USA), and dry weight (DW, mg) was obtained after 24 h of freeze-drying, using a precision balance with 1 mg resolution.

### 2.3. Otolith analysis

Larvae were placed on a microscope slide and were rehydrated with 1 or 2 drops of distilled water to facilitate otolith extraction. Sagital otoliths (hereafter, otoliths) were extracted using fine needles and cleaned with distilled water. Once dryed, otoliths were glued onto the slide with clear nail enamel (García et al., 2003). Otoliths were imaged using a Leica DM6 B microscope at 1000× magnification, with otolith metrics were measured using Leica Application Suite X (LAS X v2.0.0). Daily increments (AGE) were counted and age-corrected for larval age estimation (days) following established protocols for ABT, as detailed in Malca et al. (2023).

### 2.4. Stable isotope analysis (SIA)

Individual larvae selected for growth analysis were also processed for stable isotope analysis (SIA). Natural abundances of nitrogen (δ¹⁵N) and carbon (δ¹³C) were measured using a Thermo-Finnigan Delta-Plus isotope-ratio mass spectrometer coupled to a FlashEA1112 elemental analyzer at the Instrumental Analysis Unit of the University of A Coruña (Spain). Isotope ratios were expressed in δ notation relative to international standards: atmospheric N₂ for δ¹⁵N and Pee Dee Belemnite (PDB) for δ¹³C. Acetanilide was used as the internal standard. Analytical precision was ±0.11‰ for δ¹⁵N and ±0.14‰ for δ¹³C, based on internal reference standard deviation (duplicate repeatability).

Lipid extraction was not possible due to the limited quantity of our samples. Therefore, a subsequent lipid content correction of the δ^13^C values was conducted based on Logan et al. (2008). From the average of the three equations for all fish tissues, we applied the model best suited for whole-body samples to predict and correct for lipid content in our larval δ^13^C data. This procedure has been previously described by Laiz-Carríon et al. (2013, 2015, 2019, this issue).

### 2.5. Estimation of maternal isotopic signatures

Maternal δ-values refers to isotopic signatures inherited by offspring directly from their mother and were estimated using the model proposed for ABT by Uriarte et al. (2016) and previously utilized by Quintanilla et al. (2023, 2024):

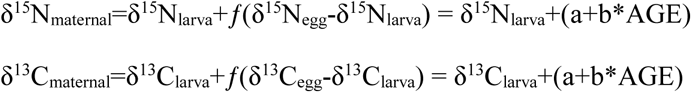

Here, δ¹⁵Nₗₐᵣᵥₐ and δ¹³Cₗₐᵣᵥₐ represent the stable isotope values of individual pre-flexion larvae, δ¹⁵Nₑ_gg_ and δ¹³Cₑ_gg_ are the average isotopic values from egg pools, and *a* and *b* are model coefficients. AGE refers to larval age (days).

The δ¹⁵N_egg_ and δ¹³C_egg_ isotopic signatures of SBT were obtained from pooled spawned eggs (n = 6) collected with plankton nets from wild samples during the IO spawning peak and were genetically identified by COI sequencing (Malca et al. this issue). ABT egg samples included in GBM were collected from a pooled batch of newly spawned eggs (n = 30) obtained from broodstock held in an open-sea cage in the western Mediterranean (Uriarte et al., 2016).

In both cases, the maternal isotopic signature assigned to each larva was simulated by randomly sampling from a normal distribution defined by the mean and standard deviation of egg isotopic values obtained from field (SBT) or experimental (GBM) data.

### 2.6. Maternal isotopic niche breadth

Maternal isotopic niches were assessed using the estimated δ¹⁵N and δ¹³C maternal values from preflexion stage larvae. Niche breadths were quantified using standard Bayesian ellipse areas corrected for small sample sizes (SEAc), with associated credible intervals (Jackson et al., 2011, 2012).

SBT and GBM populations were categorized as optimal (OPT) and deficient (DEF) growth groups according to their SL and DW residual analyses. Least squares linear regression (y = a+x*b) for log-transformed SL and log-transformed DW vs AGE were fitted to define the daily growth pattern (Figs. S1 and S2), residuals were calculated for each larval group (SBT and GBM). The two contrasting growth groups were assigned following Quintanilla et al. (2015) according to their residual values: OPT larvae had positive residuals for both variables (were larger and heavier), while DEF larvae had negative residuals for both variables (were smaller and lighter). In addition, there were two intermediate groups with mixed residuals not considered in this study. This residual-based classification identified larvae with optimal vs. deficient growth relative to the sampled bluefin tuna populations and enabled intra-population trophic comparisons between contrasting pre-flexion growth conditions.

Niche analysis was performed using the R package SIBER (v3.3.0). Bivariate standard ellipses were calculated from 40% of the variance-covariance matrix following Laiz-Carrión et al. (2019). Additionally, kernel utilization density (KUD, 40% contour) was calculated using the rKIN package (Eckrich et al., 2020), a method robust to outliers and effective for multimodal distributions.

### 2.6. Larval growth rates

The overall GBM growth rate was estimated as the slope (b) of a linear model fitted to the entire larval pre-flexion dataset (SL = a + b*AGE) (Fig 2a). Individual growth rates were calculated for each larva using the biological intercept derived from the overall GBM model (a = 3.16), based on the following equation: b = (SL – a) / AGE (Fig 2b).

**Figure 2.**
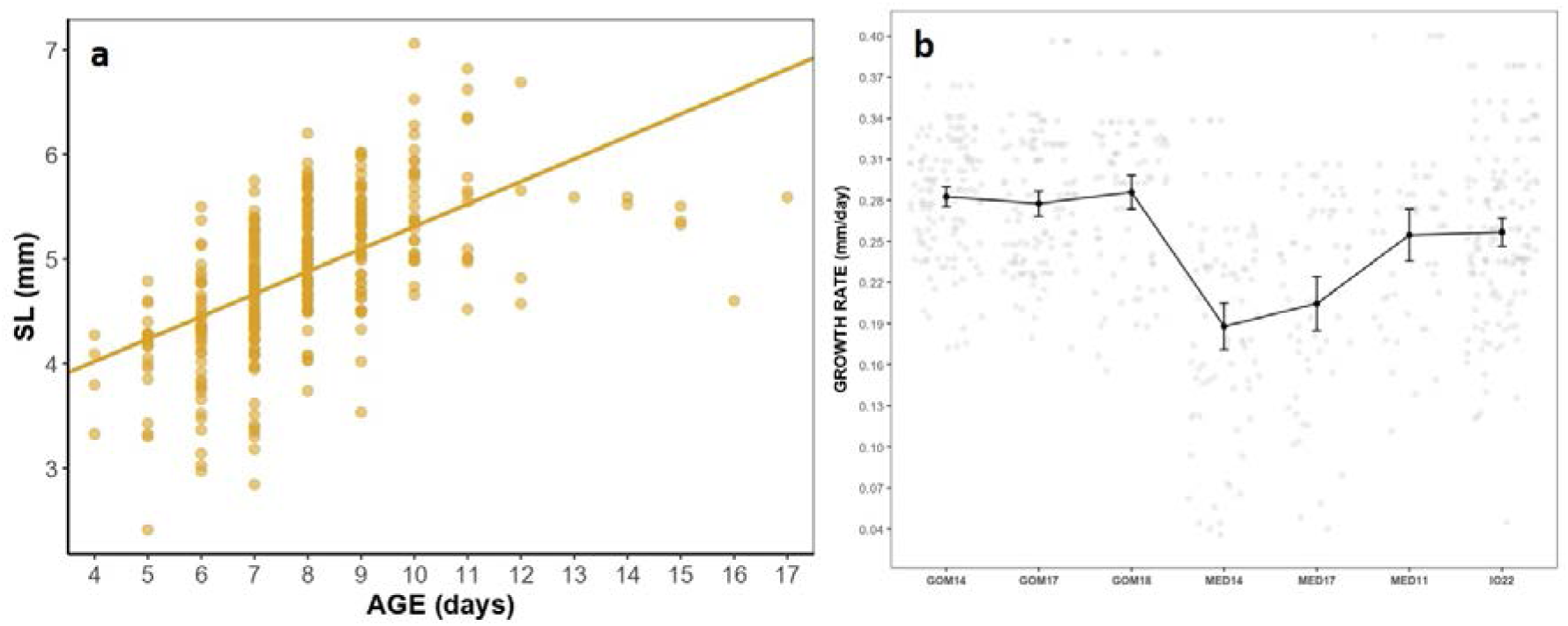
a) Relationship between standard length (SL) and age (AGE) for GBM pre-flexion bluefin tuna larvae. b) The growth rates (mm day^−1^) are indicated by the mean ± 95% confidence intervals for each population included in the GBM

### 2.7. Statistical analyses

Analysis of covariance (ANCOVA) was used to compare growth and isotopic values between OPT and DEF larval groups, using AGE as the covariate. Variables were log-transformed as necessary to meet assumptions of linearity and homoscedasticity (Sokal and Rohlf, 1979).

Estimated maternal isotopic signature values were compared between OPT and DEF using the non-parametric Mann–Whitney U test due to violation of parametric assumptions. All analyses were performed in R v4.2.1 (R Core Team, 2022) using RStudio, with significance set at α = 0.05.

## 3. Results

### 3.1. Larval growth and isotopic signatures

#### 3.1.1. Southern Bluefin Tuna

A total of 80 pre-flexion larvae from the SBT population, ranging from 2.97 to 6.34 mm SL, were analysed (Table 1). Larval δ¹⁵N and δ¹³C values showed a significant and negative linear relationship with age (Figs. 3a, b; Table 2). SIA within SBT population revealed that larvae in the positive residual group (OPT) had significantly lower δ¹⁵N (Fig. 3c, Table 3) and δ¹³C values (Fig. 3d, Table 3) compared to deficient growth group (DEF).

**Figure 3.**
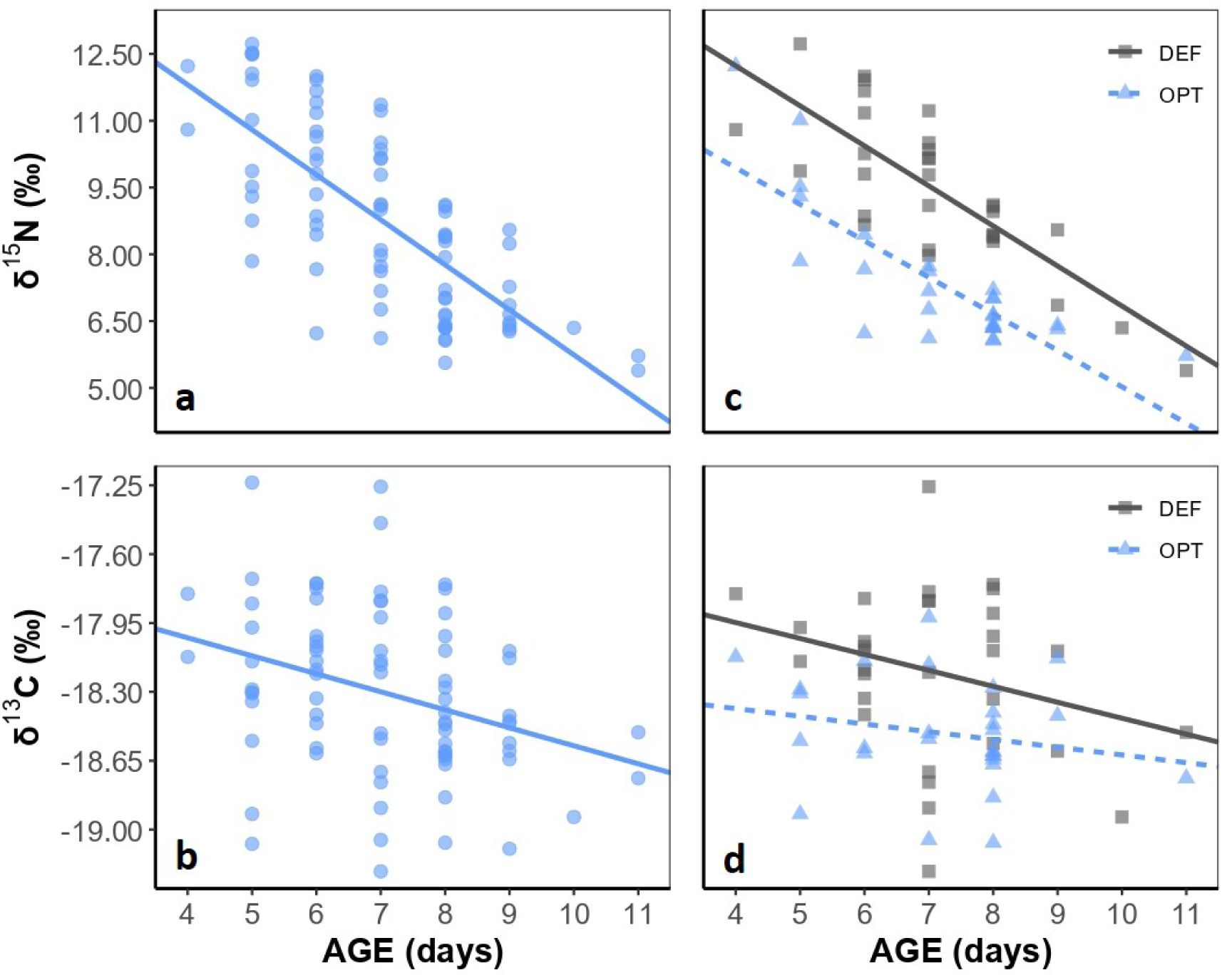
Isotopic signature of Nitrogen (a; δ^15^N) and Carbon (b; δ^13^N) with AGE (days) for the total population of SBT pre-flexion larvae 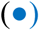 and for the two groups established by the positive and negative residuals of length and weight related to age [c and d; OPT 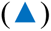 vs DEF 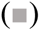].

**Table 2.** Equations for larval isotopic variables (δ¹⁵N, δ¹³C) in relation to age (AGE) for SBT and GBM. ** *p* < 0.01.

| Population | n | Isotopic variables vs AGE | *p* | $R^2$ |
| --- | --- | --- | --- | --- |
| SBT | 80 | $\delta^{15}\text{N}_{\text{larva}} = 15,85 - 1,01 * \text{AGE}$ | ** | 0,54 |
| | | $\delta^{13}\text{C}_{\text{larva}} = -17,66 - 0,09 * \text{AGE}$ | ** | 0,11 |
| GBM | 355 | $\delta^{15}\text{N}_{\text{larva}} = 10,42 - 0,53 * \text{AGE}$ | ** | 0,25 |
| | | $\delta^{13}\text{C}_{\text{larva}} = -18,64 - 0,04 * \text{AGE}$ | ** | 0,02 |

**Table 3.** Summary of Mann-Whitney U test (mean ± SE, number of larvae, Z adjusted and *p* values) within each population (SBT and GBM) in the two groups established from the positive (OPT) and negative (DEF) residuals of length (SL) and weight (DW) related to age for larval isotopic signatures and estimated maternal isotopic signatures. ** *p* < 0.01.

|  | SBT |  |  |  |  |  | GBM |  |  |  |  |  |
| --- | --- | --- | --- | --- | --- | --- | --- | --- | --- | --- | --- | --- |
|  | OPT |  | DEF |  | MW-U test |  | OPT |  | DEF |  | MW-U test |  |
| | Mean ± SE | n | Mean ± SE | n | Z-adjusted | p | Mean ± SE | n | Mean ± SE | n | Z-adjusted | p |
| $\delta^{15}\text{N}_{\text{larva}}$ | 7,31 ± 1,55 | 28 | 9,45 ± 1,67 | 31 | 4,432 | ** | 5,56 ± 1,27 | 160 | 7,56 ± 2,27 | 88 | 6,973 | ** |
| $\delta^{13}\text{C}_{\text{larva}}$ | -18,51 ± 0,28 | 28 | -18,19 ± 0,42 | 31 | 3,248 | ** | -19,07 ± 0,44 | 160 | -18,80 ± 0,68 | 88 | 2,773 | ** |
| $\delta^{15}\text{N}_{\text{maternal}}$ | 10,45 ± 2,15 | 28 | 12,55 ± 1,05 | 31 | 5,586 | ** | 11,39 ± 1,12 | 160 | 13,31 ± 1,84 | 88 | 7,878 | ** |
| $\delta^{13}\text{C}_{\text{maternal}}$ | -20,20 ± 0,28 | 28 | -19,90 ± 0,40 | 31 | 3,355 | ** | -18,82 ± 0,44 | 160 | -18,55 ± 0,69 | 88 | 2,795 | ** |

#### 3.1.2. General Bluefin Model

The GBM included 80 pre-flexion SBT larvae along with the six ABT populations (Table 1) that included 139 preflexion larvae from three GOM surveys (GOM14, GOM17, and GOM18) and 136 preflexion larvae from three MED surveys (MED11, MED14, and MED17) with a size range between 2.41 and 7.06 mm (Table 1). Similar to the SBT analysis, larval ABT δ¹⁵N and δ¹³C values showed a negative linear relationship with age (Figs. 4a, b; Table 2). Intra-population comparisons revealed that larvae in the positive residual group (OPT) had significantly lower δ¹⁵N (Fig. 4c, Table 3) and δ¹³C values (Fig. 4d, Table 3) than those in the negative residual group (DEF).

**Figure 4.**
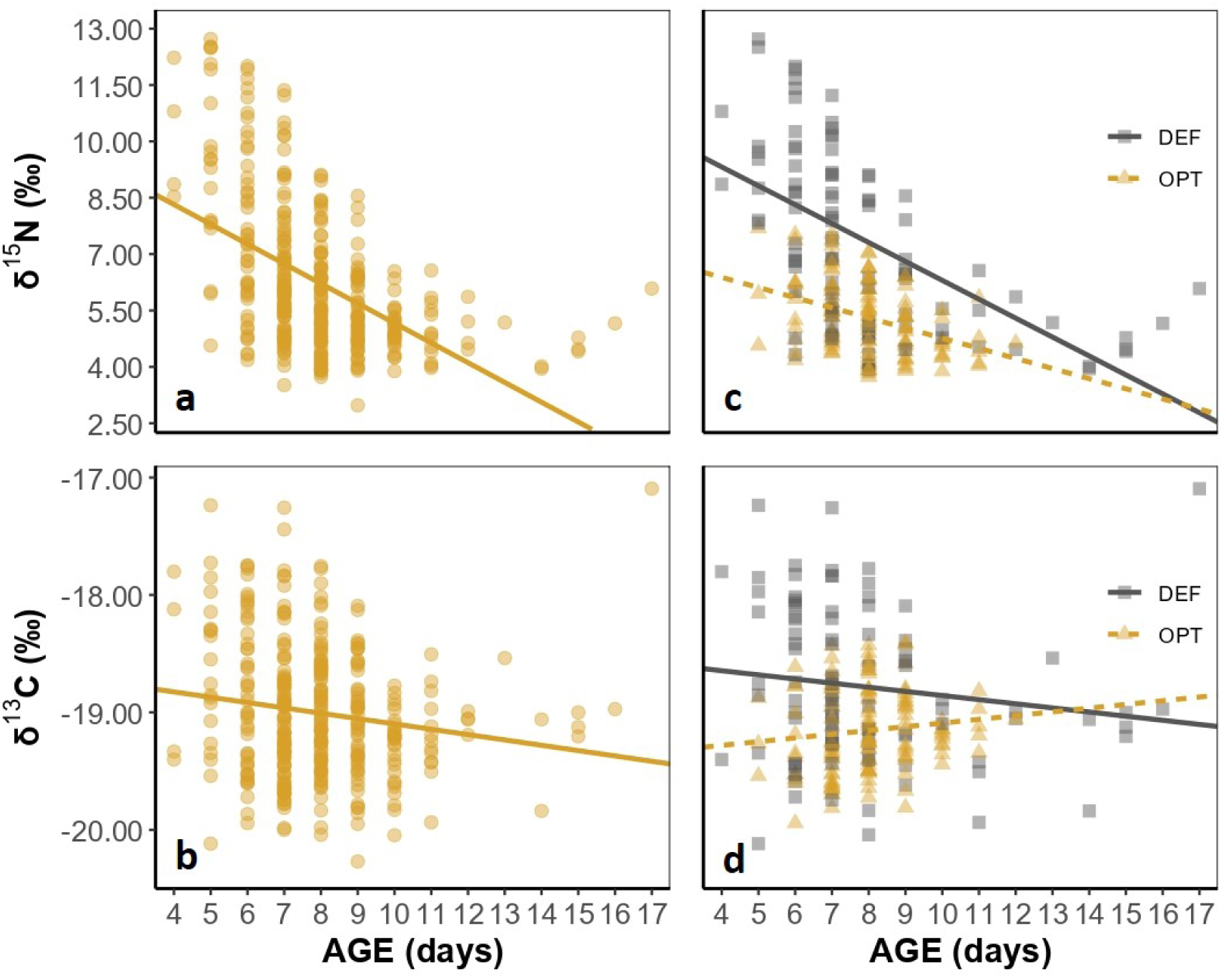
Isotopic signature of nitrogen (a; δ^15^N) and carbon (b; δ^13^C) with AGE (days) for GBM preflexion larvae 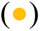 and for the two groups established by the positive and negative residuals of length and weight related to age [c and d; OPT 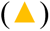 vs DEF 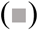].

### 3.2. Estimated maternal isotopic signatures

The equations used to calculate the age-dependent term in the ABT model proposed by Uriarte et al. (2016) applied to estimate the maternal δ-values inherited from the breeders are shown in Table 4 for both SBT and GBM. In the case of SBT, the mean ± SD values for δ¹⁵Nₑ_gg_ and δ¹³Cₑ_gg_ were 11.76 ± 0.44‰ and –19.99 ± 0.17‰, respectively. For the GBM, δ¹⁵Nₑ_gg_ and δ¹³Cₑ_gg_ had mean ± SD values of 12.10 ± 0.43‰ and –18.86 ± 0.78‰, respectively.

**Table 4.**
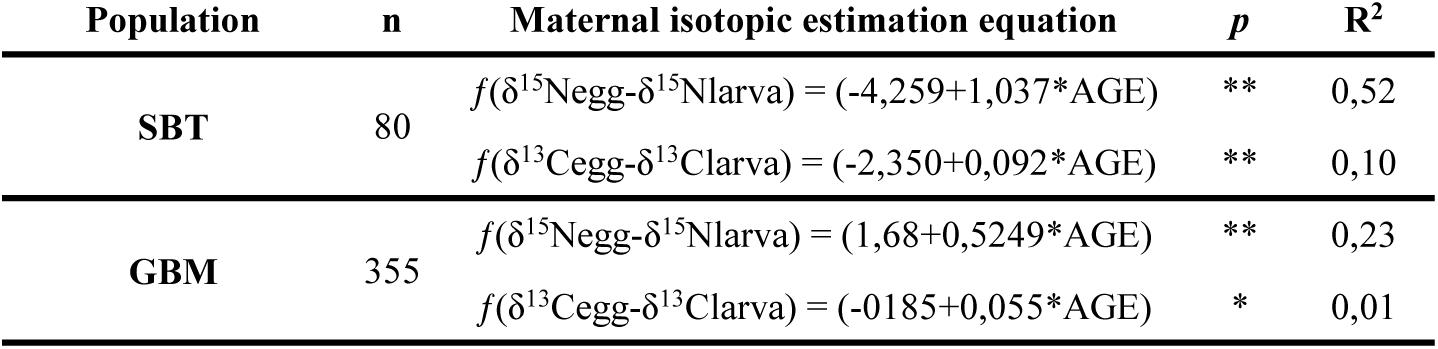
Maternal isotopic signature equations derived from pre-flexion larvae for SBT and GBM. * *p* < 0.05; ** *p* < 0.01.

| Population | n | Maternal isotopic estimation equation | p | R<sup>2</sup> |
| --- | --- | --- | --- | --- |
| SBT | 80 | $f(\delta^{15}\text{N}_{\text{egg}} - \delta^{15}\text{N}_{\text{larva}}) = (-4,259 + 1,037 * \text{AGE})$ | ** | 0,52 |
| | | $f(\delta^{13}\text{C}_{\text{egg}} - \delta^{13}\text{C}_{\text{larva}}) = (-2,350 + 0,092 * \text{AGE})$ | ** | 0,10 |
| GBM | 355 | $f(\delta^{15}\text{N}_{\text{egg}} - \delta^{15}\text{N}_{\text{larva}}) = (1,68 + 0,5249 * \text{AGE})$ | ** | 0,23 |
| | | $f(\delta^{13}\text{C}_{\text{egg}} - \delta^{13}\text{C}_{\text{larva}}) = (-0,185 + 0,055 * \text{AGE})$ | * | 0,01 |

### 3.3. Maternal isotopic niches

The comparison of residual groups showed that faster growing larvae (OPT) had averaged lower isotopic values for larval and estimated maternal values of δ^15^N and δ^13^C for SBT and GBM (Table 3). Moreover, optimal growth groups showed narrower maternal niches in both populations (Figs. 5b-c, 6b-c; Table 5).

**Figure 5.**
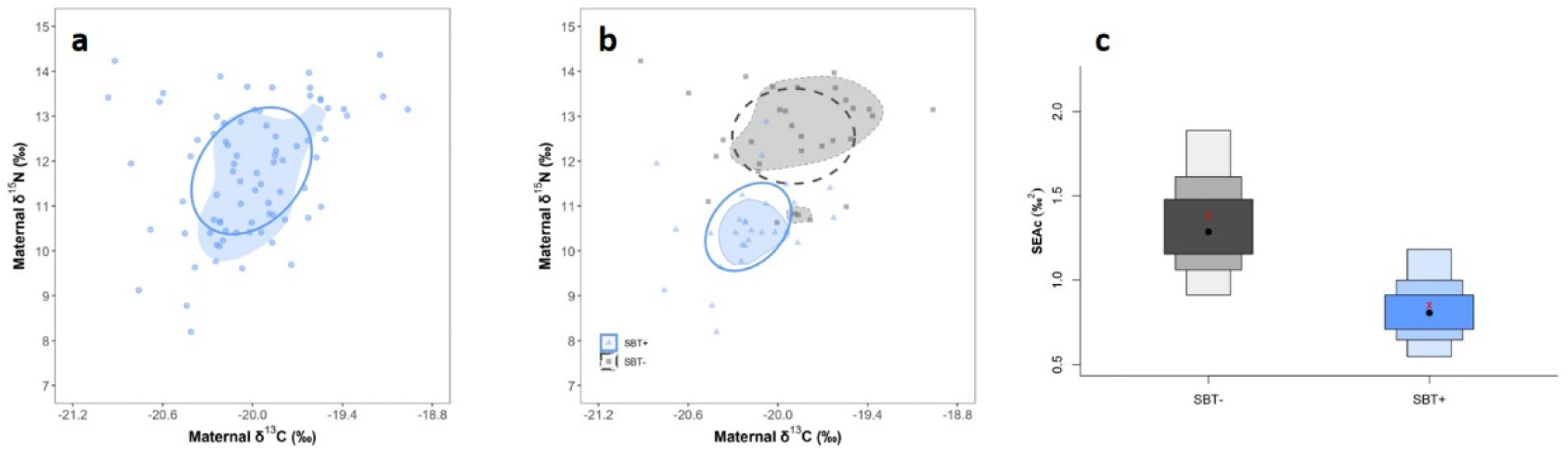
a) Biplot of δ^13^C_maternal_ vs δ^15^N_maternal_ for estimated maternal niches for SBT using SIBER (open ellipse) and rKIN (blue shaded area) packages based on pre-flexion larvae. b) Biplot of δ^13^C_maternal_ vs δ^15^N_maternal_ for estimated maternal niches for OPT (blue open ellipse; blue shaded area) and DEF (black dashed ellipse; grey shaded areas). c) Standard ellipse areas (SEAc) estimated as maternal trophic niche width for OPT and DEF using SIBER analysis. The darkest, intermediate and lightest grey boxes are the 50%, 75% and 95% credibility intervals, respectively. The red symbol (x) is the standard ellipse area calculated using correction for small sample size (SEAc).

**Table 5.** Maternal trophic niche breadth of SBT and GBM for the OPT and DEF groups, estimated using SIBER and rKIN analyses.

|  | SIBER |  | rKIN |  |
| --- | --- | --- | --- | --- |
|  | OPT | DEF | OPT | DEF |
| **SBT** | 0,85 | 1,38 | 0,49 | 1,57 |
| **GBM** | 1,03 | 3,23 | 1,36 | 1,18 |

### 3.4. Optimal maternal feeding isotopic niche (OMFIN)

GBM larvae were categorized into four groups according to their growth trajectories of standard length as a function of age (SL = 3.16 + 0.22 *AGE, R^2^=0.33, *p* < 0.05; Fig. S3): the TOTAL group (n = 355), top 50% fastest-growing larvae (n = 180), top 25% (n = 90), and top 5% (n = 20). Based on these growth rates groupings and their estimated maternal niches, Figures 7a, b and Table 6 illustrate a consistent decrease in maternal niche width with increasing larval growth rates.

**Table 6.** Maternal trophic niche breadth of GBM growth rates groups estimated using SIBER and rKIN analyses.

|  | SIBER | rKIN |
| --- | --- | --- |
| **TOTAL** | 2.46 | 2.39 |
| **50% HGRs** | 1.80 | 1.84 |
| **25% HGRs** | 1.44 | 1.67 |
| **5% HGRs** | 0.88 | 1.14 |

## 4. Discussion

Maternal effects play a crucial role in regulating offspring traits (Moore et al., 2019), significantly reducing predation and starvation risk by preventing against adverse environmental conditions and accelerating early development (Marshall et al., 2008). The magnitude of maternal influence on early larval growth is species-specific and can vary depending on the reproductive strategies, age structure of the breeding population and the nutritional quality of the maternal diet prior to reproduction.

In the subsections below, we consider the ecological link between maternal foraging and larval survival by analyzing the estimated SIA of spawning females and their influence on offspring fitness during the pre-flexion stages. We observed a faster SBT growth rates in larvae with the most constrained maternal niches and applied our findings to formulate a hypothesis for bluefin tuna species (OMFIN) developed from seven bluefin tuna populations. Considering that bluefin tuna species spawn and feed in separate areas, during different seasons, by storing energy and drawing on it later after long-distance migrations to spawning grounds, our hypothesis offers insights into maternal trophic ecology of bluefin tunas

### 4.1. Stable isotope patterns in SBT larvae

The decrease in δ¹⁵N values observed in pre-flexion larvae of SBT with age (Fig. 3a; Table 2) are consistent with results from larval ABT rearing experiments (Uriarte et al., 2016) and field studies for the same species (García et al., 2017; Laiz-Carrión et al., 2019; Quintanilla et al., 2023, 2024). These studies suggest that high δ¹⁵N values in early larval stages result from maternal isotope transfer, which progressively decreases with ontogeny. These δ¹⁵N differences between pre- and post-flexion stages have been linked with energy acquisition and allocation to egg production and interpreted as a capital breading behaviour when a fish species spawn and feed in separate areas, during different seasons by storing energy and utilizing it later for reproduction (McBride et al., 2015; Mei et al., 2018; Laiz-Carrión et al., 2019)

In our intra-population growth classification, SBT larvae exhibited distinct linear relationships between δ¹⁵N and age, with significantly lower δ¹⁵N values in faster-growing individuals. This ontogenic pattern mirrors previous observations in ABT larvae (Quintanilla et al., 2023, 2024). Three main hypotheses have been proposed to explain the link between lower δ¹⁵N values and enhanced larval growth: (1) differences in breeder age, (2) variations in maternal nutritional condition, and (3) batch-specific variability in egg quality (Quintanilla et al., 2023). On the other hand, the lower δ¹³C values observed in larvae from the OPT group suggest that the origin of the food sources of breeders also plays a key role in SBT larval growth

### 4.2. Maternal isotopic niche and larval growth in SBT

Maternal food sources are a key determinant of egg and larvae biochemical composition (Yoshida et al., 2011; Schlotz et al., 2013, Planas, 2021, Planas et al. 2021). Intraspecific dietary variation (Bolnick et al., 2002; Griffen, 2014) can result in differences in egg size and energy reserves, directly impacting larval growth. Isotopic niche breadth is inherently plastic and can vary in response to resource availability (Lesser et al., 2020), often along productivity gradients (MacArthur and Pianka, 1966; O’Farrell et al., 2014). In productive environments, niche breadth tends to narrow as abundant resources allow for specialized diets. Conversely, under low productivity, individuals may adopt broader foraging strategies to meet metabolic demands, resulting in more diverse diets with wider isotopic niches (Lesser et al., 2020). Within this framework, our results for SBT larvae suggest that optimal growing individuals are linked to more specialized (stenophagous) maternal feeding strategies, reflected in narrower maternal isotopic niches (Fig. 5b, c; Table 5). This association has been previously reported for ABT (Quintanilla et al., 2023, 2024), supporting the idea of a direct link between larval growth potential and maternal foraging behaviour. Moreover, the differences in the mean δ¹⁵N and δ¹³C values of maternal estimates (Table 3), together with the lack of overlap between the maternal isotopic niches of the OPT and DEF groups, suggest that breeders from each group exploit distinct trophic niches with different isotopic characteristics.

### 4.3. Implications of the General Bluefin Model (GBM)

The GBM consistently revealed that faster-growing larvae are linked to more specialized maternal diets. The patterns observed in the General Bluefin Model (GBM) are consistent with those from both SBT and ABT populations. Specifically, a decline in δ¹⁵N with ontogeny (Fig. 4a, Table 2), lower δ¹⁵N values in optimally growing larvae (Fig. 4c; Table 3), and narrower maternal isotopic niches (Figs. 6b, c, Table 6) are common features associated with enhanced larval growth across bluefin species. As observed in the SBT population, the OPT group also exhibited lower δ¹³C values (Fig. 4d; Table 3), further reinforcing the role of prey sources in bluefin larval growth.

**Figure 6.**
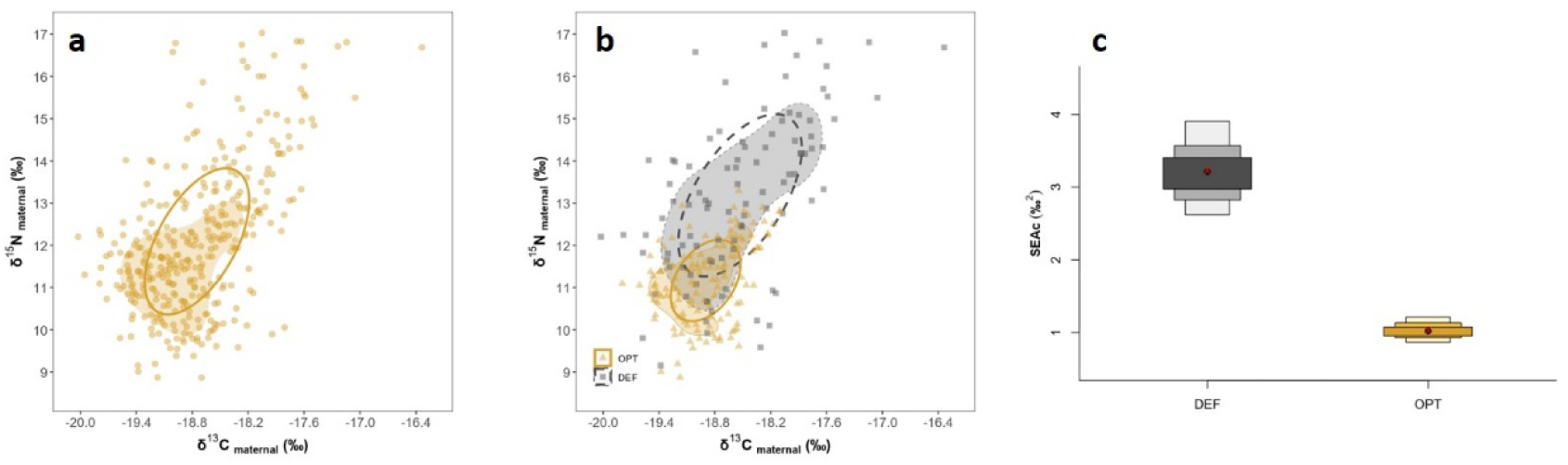
a) Biplot of δ^13^C_maternal_ vs δ^15^N_maternal_ for estimated maternal niches for the General Bluefin Model (GBM) using SIBER (open ellipse) and rKIN (shaded area) packages based on pre-flexion larvae. b) Biplot of δ^13^C_maternal_ vs δ^15^N_maternal_ for estimated maternal niches for OPT (open yellow ellipse; yellow shaded area) and DEF (dashed black ellipse; grey shaded area). c) Standard ellipse area estimated as maternal trophic niche width for OPT and DEF by SIBER analysis. The darkest, intermediate and lightest grey boxes are the 50%, 75% and 95% credibility intervals, respectively. The red symbol (x) is the standard ellipse area calculated using correction for small sample size (SEAc).

**Figure 7.**
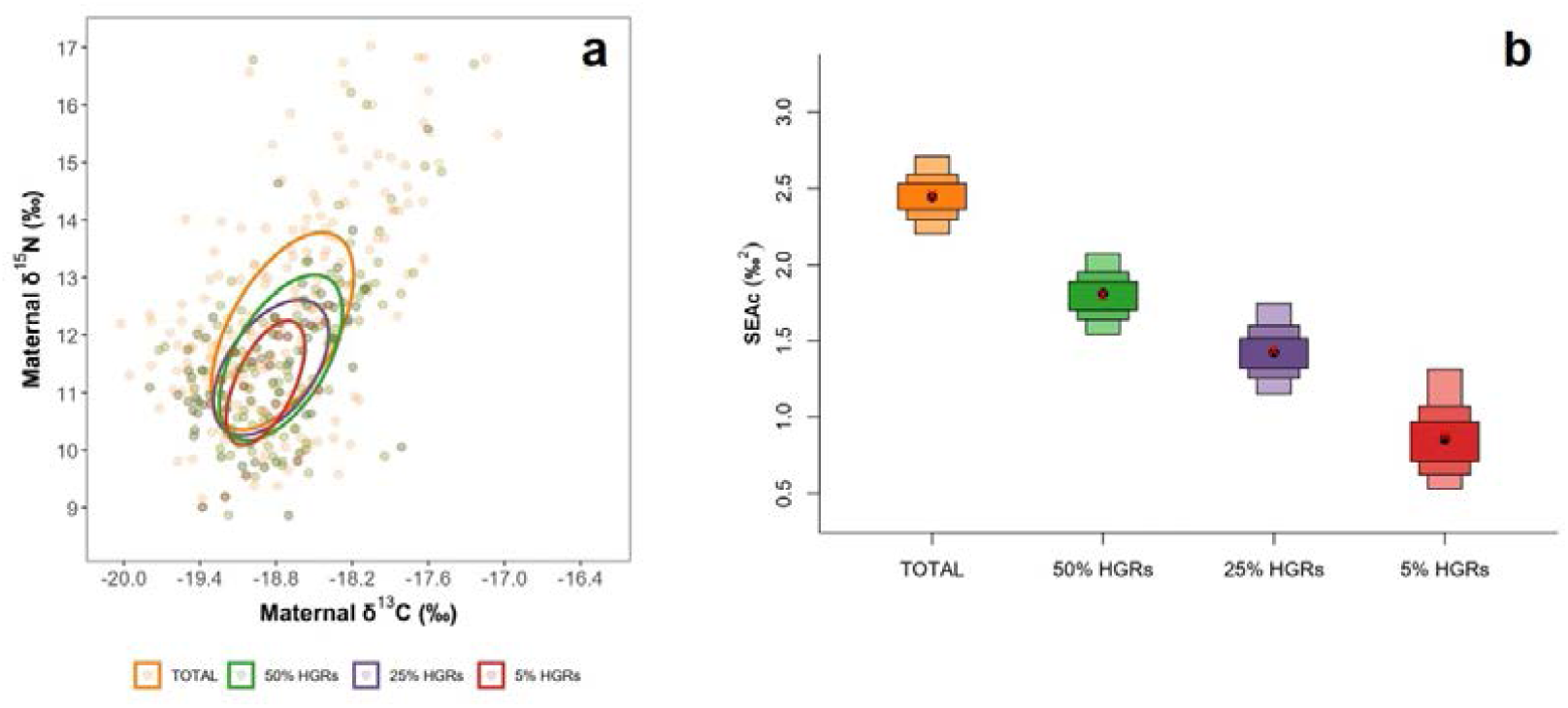
a) Biplot of δ^13^C_maternal_ vs δ^15^N_maternal_ for estimated maternal niches for the four groups (TOTAL, 50% HGRs, 25% HGRs and 5% HGRs) estimated by models based on pre-flexion larvae. b) Standard ellipse area estimated as maternal trophic niche width for every group by SIBER analysis. The dark, intermediate and light boxes are the 50%, 75% and 95% credibility intervals, respectively. The red symbol (x) is the standard ellipse area calculated using correction for small sample size (SEAc).

Remarkably, this pattern holds despite the inclusion of larvae spanning diverse years, spawning areas, growth rates, and even species. This suggests that stenophagous maternal foraging strategies may promote larval growth potential across bluefin tunas from early development stages.

Higher growth rates during early ontogeny improve survival by enhancing predator evasion, feeding efficiency, and environmental adaptability (Chick et al., 2000; Pitchford and Brindley, 2001; Takasuka et al., 2003). In SBT, this link is supported by otolith increment analyses in larvae collected within the same water mass and similar hatch dates but sampled at different time points in a survival analysis (Borrego-Santos et al., this issue).

To further explore the link between larval growth and breeder trophic patterns, we analysed maternal niche breadth across four GBM groups defined by larval growth rates (Fig. S3). While the most selective groups were dominated by fast-growing populations from the GOM (> 0.25 mm day^−1^: GOM14, GOM17, GOM18), individuals from other populations were still represented. Notably, even the most restrictive groups included individuals from MED14, the population with the lowest overall growth rate (< 0.19 mm day^−1^) (Fig 2b), highlighting the intrinsic variability in larval performance.

Maternal isotopic niches can be interpreted as trophic niches, directly reflecting the dietary sources used by breeding females. These niches allow us to infer differences in maternal diets and offer quantitative insights, such as niche width and overlap, by enabling comparisons of trophic ecology both within and between populations (Quintanilla et al. 2023). We observed that maternal niche breadth consistlenly narrowed with larval growth rates increase, with the fastest growing larvae group (5% HGRs, n = 20) exhibiting the narrowest maternal niche (Fig. 7a,b). These findings lead us to propose an Optimal Maternal Feeding Isotopic Niche (OMFIN), characterized by a restricted isotopic signature. Females foraging within it would produce larvae with superior growth rates and, by extension, increased survival potential.

According to Optimal Foraging Theory (OFT) in marine fish, individual diets variation is influenced by both the availability of prey and the individual’s phenotype, or “individual state,” such as size, sex, or developmental stage (Schoener, 1971; Werner and Hall, 1974; Svanbäck and Bolnick, 2005). Although individuals may be capable of consuming a broad range of prey types, they often select specific prey items based on the trade-off between energetic gain and the cost of prey handling (Cachera et al., 2017).

The Optimal Maternal Feeding Isotopic Niche (OMFIN) was established based on isotopic estimates of maternal niches. These estimates originated from pre-flexion larvae across seven bluefin tuna populations collected in different years and spawning regions. Even with the model’s extensive geographical, temporal and environmental variability, the constricted range of maternal δ¹⁵N and δ¹³C values linked to the fastest-growing larvae result in a compact niche definition. In this sense, the OMFIN likely reflects the ideal isotopic pattern of foraging areas that enable females to endow their offspring with higher growth potential via maternal inheritance. Our results suggest that breeders producing larvae with higher growth potential specialize on prey with consistent isotopic signatures across populations. This association between narrower niches and higher larval growth rates in the GBM reinforces the link between larval performance and breeder trophic behaviour in bluefin tunas.

## 5. Conclusions and future perspectives

Our study illuminates a crucial relationship: mothers that produce faster-growing larvae tend to specialize their diets, resulting in a narrower “Optimal Maternal Feeding Isotopic Niche” (OMFIN). These findings strengthen our conceptual framework, which posits that specialized maternal diets are linked to enhanced larval performance. This framework appears robust and the underlying mechanisms of the OMFIN may be generalizable across bluefin tuna species. The observed connection between maternal diet specialization and larval growth has significant implications for understanding reproductive success, larval survival, and recruitment in highly migratory pelagic species.

To accurately estimate maternal isotope values in future research, we’ll need species-specific ontogenetic models that describe how isotopic signatures develop. We also need data to quantify maternal transfer under various pre-spawning diets. Since baseline δ^15^N and δ^13^C values for maternal tissues are often unavailable, trophic-position estimates for the females themselves remain uncertain. Applying amino-acid compound-specific isotope analysis (AA-CSIA) could partially overcome this limitation, allowing for a more precise resolution of maternal trophic position and a better assessment of its potential influence on larval growth.

A logical next step would be to examine Pacific Bluefin Tuna (*Thunnus orientalis*) as a case study to determine if the OMFIN hypothesis also holds in the Pacific Ocean. More broadly, this hypothesis can be tested in any species where oogenic energy is acquired at times and places different from the spawning ground (i.e., capital-breeding species), by exploring how maternal diet influences larval growth and survival. Controlled feeding trials with broodstock held under experimental conditions would be particularly valuable for validating OMFIN and interpreting ecological outcomes. Species that are more amenable to captivity and easier to sample during reproduction could provide additional insight into the link between maternal nutrition, larval viability, and recruitment.

## Data Availability Statement

The data that support the findings of the present study are available from the first/corresponding author, J. M. Quintanilla, upon reasonable request

## Credit Authorship Contribution Statement

**JMQ**: Conceptualization, Formal analysis, Investigation, Methodology, Visualization, Writing-original draft, review and editing. **RBS**: Conceptualization, Formal analysis, Visualization, Writing-review and editing. **EM**: Investigation, Methodology, Visualization, Writing-review and editing. **IR**: Writing-review and editing. **FJA**: Formal analysis, Writing-review and editing. **MP**: Writing-review and editing. **RS**: Investigation, Writing-review and editing. **MRL**: Funding acquisition, Supervision, Project administration, Writing-review and editing. **RLC**: Conceptualization, Formal analysis, Funding acquisition, Investigation, Methodology, Supervision, Visualization, Project administration, Writing-original draft, review and editing.

## Declaration of Generative AI

Generative AI and AI-assisted technologies were solely employed during the writing process to improve the clarity and language quality of the manuscript.

## Acknowledgements

We thank the crew of the research ships R/V Gordon Gunter, R/V Roger Revelle, R/V Cornide de Saavedra and R/V SOCIB, for their support in conducting the corresponding surveys. We deeply thank Drs. Alberto García, John Lamkin and Trika Gerard for their contributions to establish a collaborative framework from Spain and USA.

## Supplementary material

**Figure S1.**
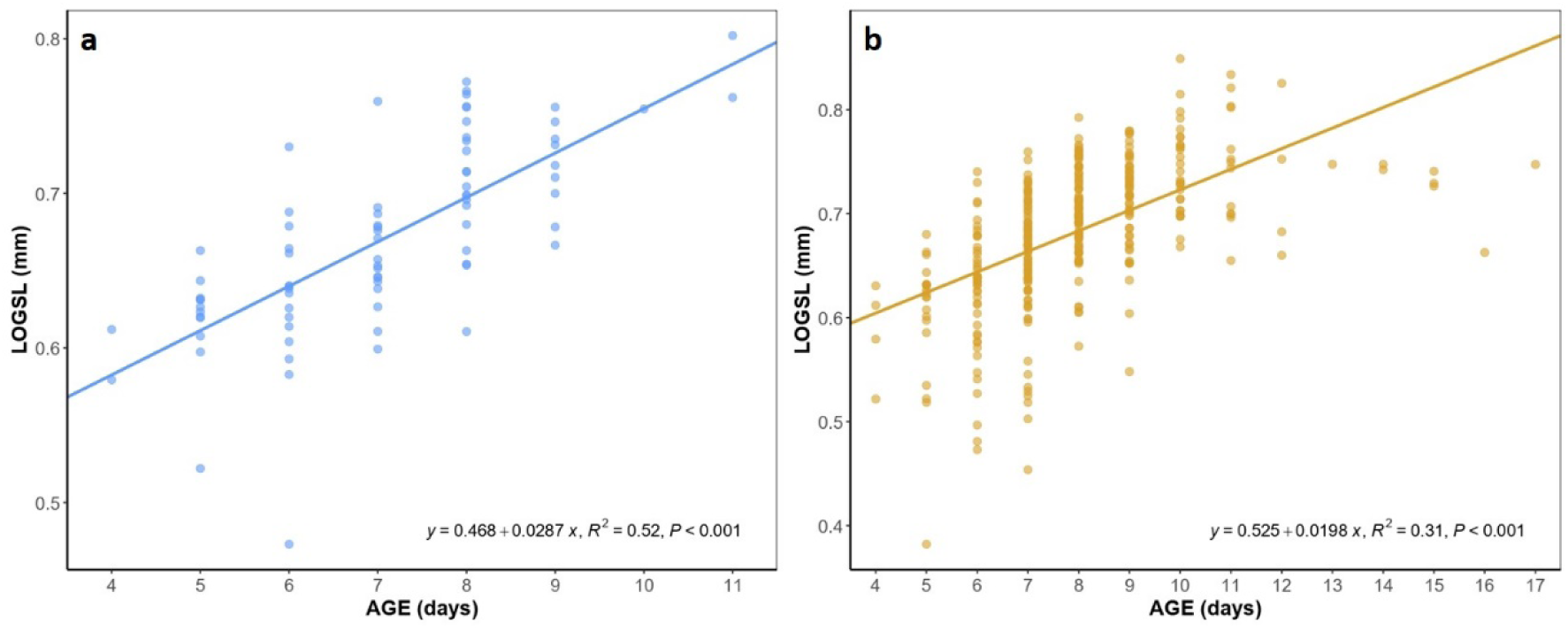
Relationship between log-transformed SL (LOGSL) and age (AGE) for SBT (a, 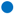) and GBM (b, 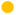) pre-flexion bluefin tuna larvae.

**Figure S2.**
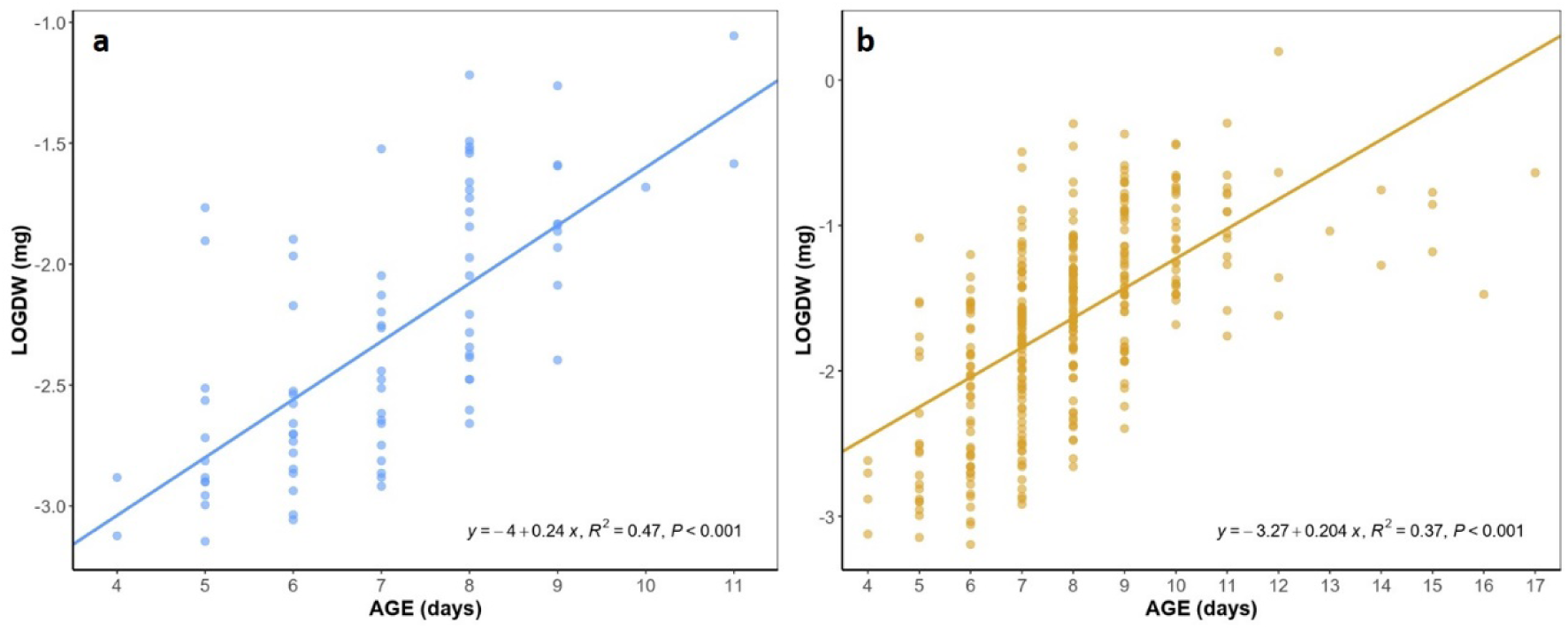
Relationship between log-transformed DW (LOGDW) and age (AGE) for SBT (a, 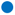) and GBM (b, 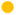) pre-flexion bluefin tuna larvae

**Figure S3.**
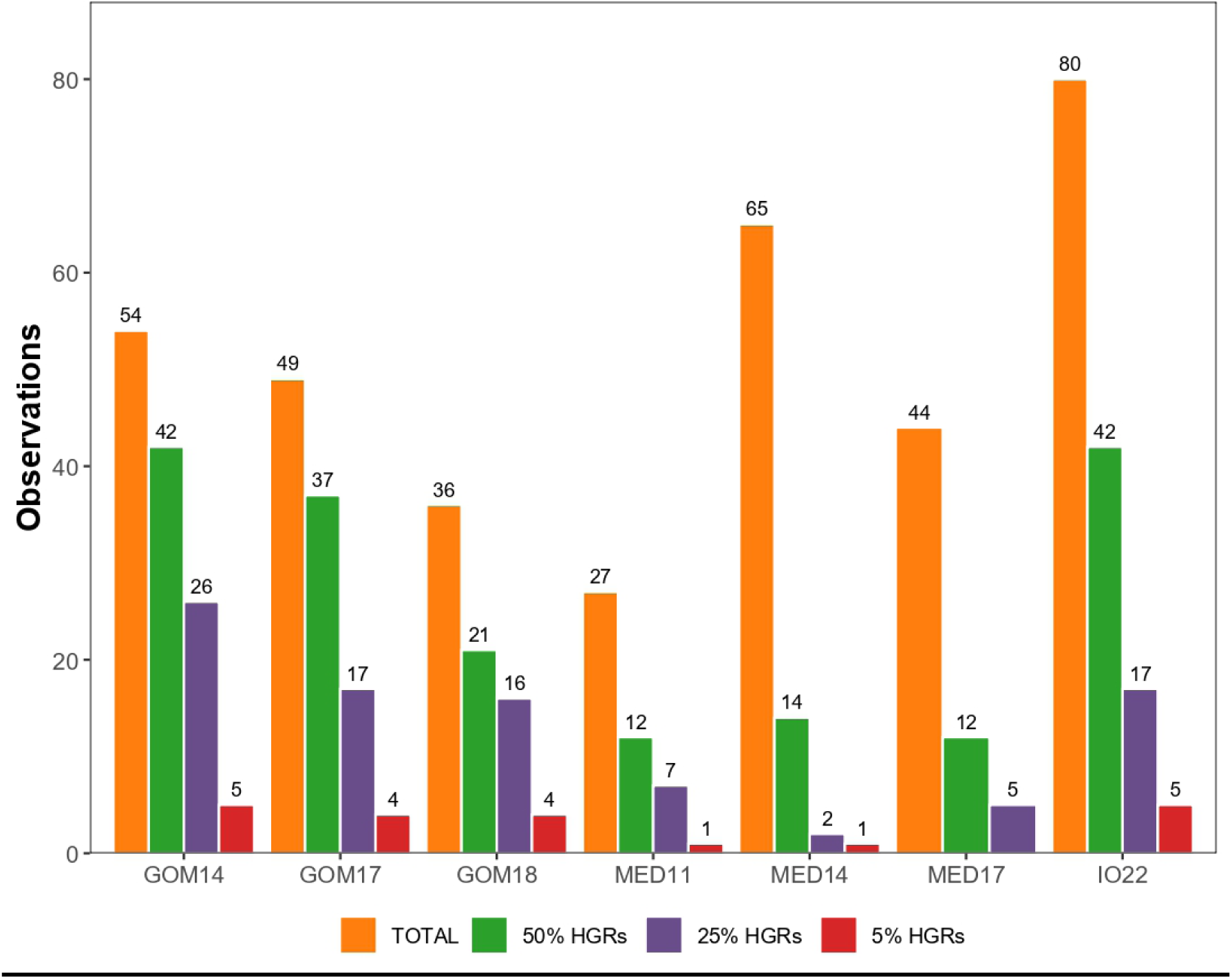
Seven bluefin tuna populations (GBM pre-flexion larvae) according to their growth rates (mm day^−1^): TOTAL (n = 355), the top 50% fastest-growing larvae (50% HGRs; n = 180), top 25% (25% HGRs; n = 90), and top 5% (5% HGRs; n = 20).

